# Microfluidic Transfection for High-Throughput Mammalian Protein Expression

**DOI:** 10.1101/200261

**Authors:** Kristina Woodruff, Sebastian J. Maerkl

**Author notes:** Correspondence should be addressed to S.J.M.

## Abstract

Mammalian synthetic biology and cell biology would greatly benefit from improved methods for highly parallel transfection, culturing and interrogation of mammalian cells. Transfection is routinely performed on high-throughput microarrays, but this setup requires manual cell culturing and precludes precise control over the cell environment. As an alternative, microfluidic transfection devices streamline cell loading and culturing. Up to 280 transfections can be implemented on the chip at high efficiency. The culturing environment is tightly regulated and chambers physically separate the transfection reactions, preventing cross-contamination. Unlike typical biological assays that rely on end-point measurements, the microfluidic chip can be integrated with high-content imaging, enabling the evaluation of cellular behavior and protein expression dynamics over time.

## SECTION 1 INTRODUCTION

Cell-based research and therapeutic endeavors frequently entail expressing and studying specific proteins. Transfection is the process of introducing foreign genetic material into mammalian cells [1]. In contrast to those produced in prokaryotic systems, the expressed proteins undergo proper folding and post-translational modifications. This technology is thus pertinent to protein production, functional assays, and therapeutic gene delivery.

Chemical transfection can be achieved with widely used reagents such as cationic lipids, which neutralize the negative charge of DNA and facilitate entry into the cell by endocytosis. The integration of lipid-based transfection with contact spotting has enabled high-throughput transfection. In this technique, called reverse transfection, purified cDNA samples are mixed with transfection reagent and spotted onto glass slides [2]. The arrays are next seeded with cells, which undergo transfection *in situ*, resulting in exogenous gene expression. Unlike protein microarrays [3–5], this method does not require individual purification of each sample and proteins can be analyzed in the natural cellular context. More than 5,000 samples can be printed on a single glass microscope slide using standard techniques [2, 6, 7]. This throughput enables massively parallel characterization of complex synthetic networks and the screening of genome-wide RNAi and cDNA libraries [8, 9].

Although reverse transfection has been optimized for a variety of genetic materials [10, 11] and cell types [12], methods for cell manipulation on the arrays still stand to be improved. Cell seeding and culturing is performed manually, and cross-contamination is a concern because spots on the array are not physically separated from one another. Standard reverse transfection offers poor control over the cell microenvironment and cannot support sophisticated downstream experiments.

Microfluidics could compensate for these shortcomings by providing a means to enclose each position on the array in a cell culture chamber. Microfluidic devices are readily fabricated using standard photolithography and soft lithography techniques [13] and contain micromechanical components such as valves that execute complex fluidic manipulations. These devices are characterized by high throughput, automation, small sample requirements, and compatibility with other analytical techniques. Standard microfluidic chips designed for mammalian cells are capable of a variety of functions, including long-term perfusion culture and the ability to individually address cell chambers [14].

This protocol describes the high-throughput transfection of mammalian cells on a microfluidic chip [15]. The device consists of two components: a glass slide patterned with transfection reagent, and a microfluidic device with chambers that support cell culturing (Fig. 1). The 280 microfluidic chambers have a capacity of ∼600 cells each. The chambers of the chip are aligned to the transfection array so that each chamber contains a unique transfection mixture. The arraying of DNA-containing transfection mixtures is coupled to the arraying of poly-L-lysine (PLL), which serves to anchor the DNA to the glass slide. This process has been optimized to enable highly efficient transfection with low cross-contamination. The chip is operated and cultured in a microscopy setup, facilitating the interrogation of protein expression dynamics and synthetic gene network performance.

**Figure 1.**
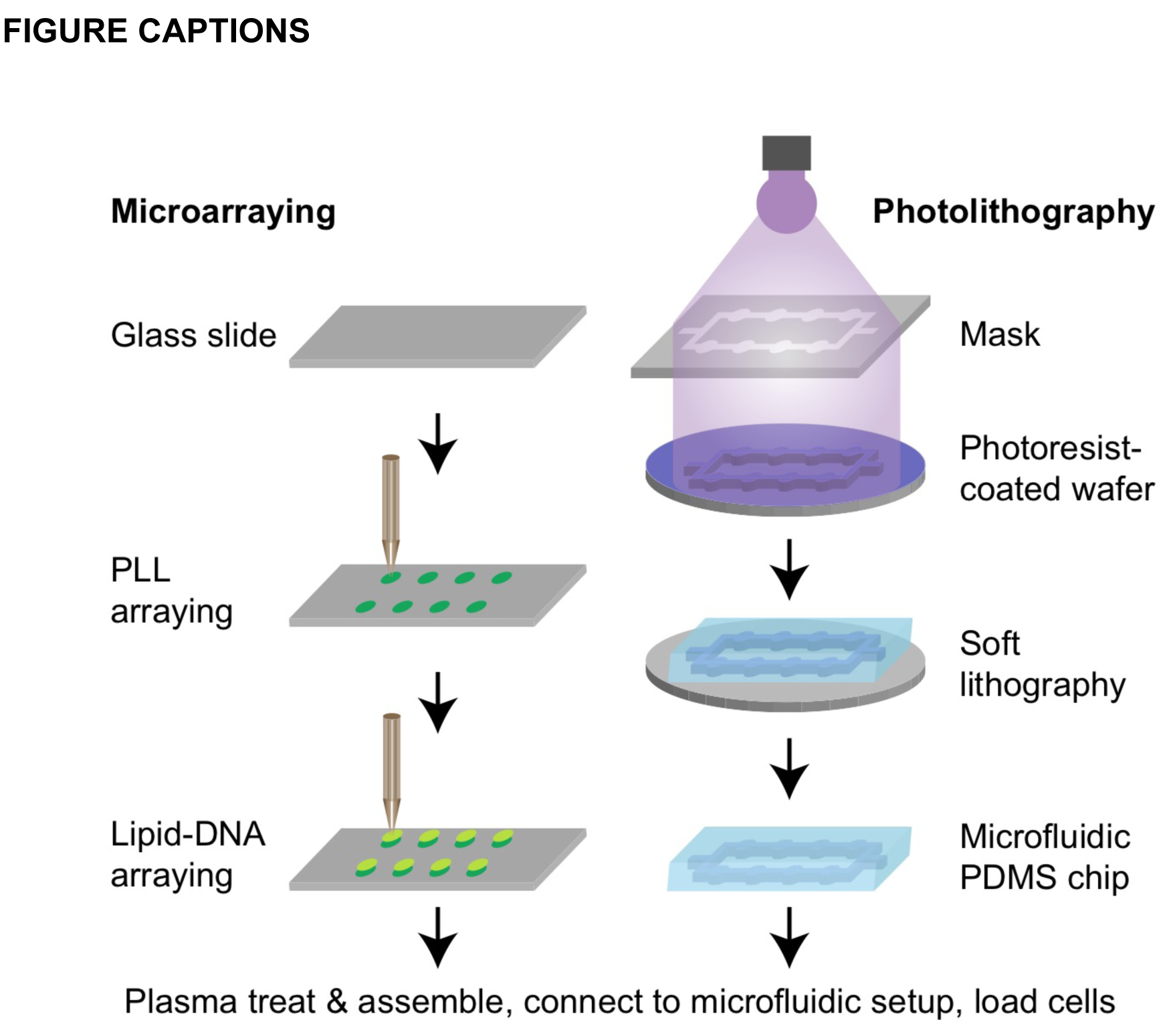
Workflow to generate a microfluidic transfection array. The transfection device consists of two components: a glass slide patterned with transfection reagent (left) and a microfluidic device that supports cell culturing (right). The lipid-DNA transfection mixtures are arrayed into distinct spots on the glass slide. The microfluidic component is fabricated using standard photolithography and soft lithography techniques. The two components are assembled together so that one lipid-DNA spot is enclosed inside each microfluidic cell culturing chamber. The assembly is connected to a microfluidic setup, and cells loaded into the device undergo transfection.

## SECTION 2 MATERIALS

### 2.1. Mask and wafer fabrication

#### 2.1.1. Cleanroom equipment for photolithography

- Computer-aided design (CAD) software.
- Heidelberg VPG200 laser lithography system (Heidelberg Instruments Mikrotechnik GmbH).
- DV10 developer (Süss MicroTec AG).
- Tepla300 dry etcher, oxygen plasma (PVA Tepla AG).
- MA6 mask aligner (Süss MicroTec AG).
- Sawatec LSM250 programmable coater for SU-8 (Sawatec AG).
- EVG150 coater and developer system for positive resist (EV Group).
- Dektak XT surface profilometer (Bruker).

#### 2.1.2. Chemicals and materials for photolithography

- Silicon wafers, diameter: 100 ± 0.5 mm, thickness: 525 ± 25 μm, conductivity type: N or P, dopant: Boron or Phosphorous, resistivity range: 0.1–100 Ωcm (Okmetic).
- Chrome-coated mask plates, 5x5 inches (Nanofilm).
- Developer MP 351 (Merck).
- Chromium etch CR7 MOS: (NH_4_)_2_Ce(NO_3_)_6_; HClO_4_.
- TechniStrip P1316 (Microchemicals).
- AZ9260 photoresist (Clariant GmbH).
- Developer AZ 400K (Merck).
- GM1070 (Gersteltec).
- Propylene glycol methyl ether acetate (PGMEA).
- Isopropanol.

### 2.2. PDMS chip fabrication

#### 2.2.1. Equipment for soft lithography

- Thinky mixer ARE-250 with adaptor for 100 ml PP beakers (C3 Prozessund Analysentechnik GmbH).
- Vacuum desiccator (Fisher Scientific AG).
- SCS G3P-8 spin coater (Specialty Coating Systems).
- Manual hole punching machine and pin vises, 21 gauge, 0.04” OD (Technical Innovations, Inc.).
- Stereomicroscope SMZ1500 (Nikon AG).
- Femto plasma cleaner (Diener electronic).
- 80°C oven.

#### 2.2.2. Materials for soft lithography

- Trimethylchlorosilane (TMCS) (Sigma).
- Polydimethylsiloxane (PDMS), Sylgard 184 (Dow Corning).
- 100 ml polypropylene (PP) beakers.
- 10-cm diameter glass Petri dishes.
- Tweezers, scalpel.
- Scotch tape.
- Aluminum foil.

### 2.3. Microarray fabrication

- QArray2 microarrayer (Genetix GmbH).
- Conical-well, poly(propylene) 384-well plates (Arrayit).
- Arraying pin with a 300 μm spot diameter and 3.3 nl delivery volume (946MP9, Arrayit).
- Standard glass microscope slides, 76 x 26 x 1 mm (VWR).
- Glass slide rack and dish.
- Dish soap.

### 2.4. PLL and transfection mixture preparation

- NaOH.
- Ethanol.
- Milli-Q water.
- 0.1% poly-l-lysine (PLL) solution (Sigma).
- Boric acid (Sigma).
- Sucrose (Sigma).
- Gelatin, Type B, 225 g Bloom (G-9391, Sigma).
- 0.1% fibronectin solution (F0895, Sigma) or bovine plasma fibronectin (F4759, Sigma, dissolved in water at 1 mg/ml).
- Supercoiled plasmid DNA (see Note 1); eGFP and tdTomato with CMV promoters.
- Effectene (Qiagen).
- 20 ml syringe with Luer-Lok tip.
- 25 mm syringe filter with 0.45 μm cellulose acetate membrane.
- Vortex.
- Small bench-top centrifuge.

### 2.5. Cell culture

- HEK 293-T cells.
- Autoclave.
- Dulbecco’s Modified Eagle Medium (DMEM) (Life Technologies).
- Fetal bovine serum (FBS) (see Note 2) (Life Technologies).
- Antibiotic-antimycotic (Life Technologies).
- TrypleE express (Life Technologies).
- CO_2_ independent medium (18045054, Life Technologies).
- GlutaMAX (Life Technologies).
- T-75 flasks, serological pipets, 0.2 μm filter, other standard cell culture materials.
- 25 ml laboratory bottles (Duran) with open-top cap (GL25, Schott) and silicone septa.

### 2.6. Microfluidic setup [16]

- Precision pressure regulator, BelloFram Type 10, 2–25 psi, 1/8” port size (960-001-000, Bachofen SA).
- Bourdon tube pressure gauges, 0–30 psi, G 1/4 male connection (NG 63-RD23-B4, Kobold Instruments AG).
- Custom-designed manual manifolds (rectangular metal casing: 14.5 x 1 x 1”) with toggles and barbs for 1/16” ID tubing, 1/4 NPT connection (Pneumadyne Inc.).
- Fittings to connect regulators to gauges and to luer manifolds: tee union (SO 03021-8) and male adaptor union (SO 01121-8-1/8) (Serto AG).
- Polycarbonate luer fittings (Fisher Scientific AG), multiport luer manifolds for flow inlet regulation (06464-87, Cole Parmer); male luer to luer connector (06464-90, Cole Parmer).
- Tygon tubing for pneumatic setup, 1/4” OD x 1/8” ID (Fisher Scientific AG).
- Disposable stainless steel dispensing needles to connect to syringe, 23 gauge, 1/2” long, 0.33 mm ID (560014, I and Peter Gonano).
- Flexible plastic tubing for fluidic connections, Tygon S54HL, 0.51 mm ID (Fisher Scientific AG).
- PTFE tubing for fluidic connections, 0.022" ID x 0.042" OD (Cole Parmer).
- Steel pins for chip-to-tube interface, Tube AISI 304 OD/ID x L 0.65/0.30 x 8 mm, cut, deburred, passivated (Unimed SA).

### 2.7. Microscope setup

- Nikon Ti-E Eclipse automated epi-fluorescence microscope.
- Filter cubes: TexasRed (HC 562/40, HC 624/40, BS 593) and FITC (HC 482/35, HC 536/40, BS 506) (AHF Analysentechnik AG).
- Cube air heater and incubation chamber to enclose the microscope (Life Imaging Services).
- Ixon DU-888 camera (Andor Technology).
- NIS Elements, Fiji, and MATLAB imaging and image processing software.

## SECTION 3 METHODS

### Methods overview

High-throughput microfluidic transfection incorporates several techniques including photolithography, soft lithography, microarraying, operation of a microfluidic setup, and cell culture (Fig. 1). The 280-chamber transfection device consists of 2 layers: a flow layer (containing cells, media, etc.) and a superimposed control layer (valves to manipulate the flow layer). Both layers are written on chrome masks in a clean room facility. The masks transfer the design to photoresist-coated wafers, which serve as molds for PDMS casting. Soft lithography techniques are employed to fabricate the PDMS chips.

The PDMS chip is aligned and bonded to a transfection microarray to generate the final device. The microarray is similar to those used for reverse transfection [2, 17– 19] and consists of 280 spots containing DNA mixed with a lipid-based transfection reagent. The surface of the microarray is patterned with PLL (a hydrophilic, DNA-retaining substance) in the positions of the chambers prior to lipid-DNA microarraying (Fig. 1). This protocol ensures robust DNA attachment, even under the high flow rates experienced on the microfluidic chip. Unlike traditional approaches that use glass slides entirely covered in PLL, this method is compatible with hydrophobic PDMS chips since the chip is not directly in contact with the positions coated with PLL.

The assembled device is connected to a microfluidic setup. A suspension of cells is loaded into the channels, and valves are actuated to segregate individual chambers. Medium is perfused through channels running parallel to the cell chambers, eliminating sheer stress during culturing. Diffusion from the channels introduces nutrients into and eliminates waste from the chambers. The cell chip is cultured directly on a microscope stage encased in an incubation chamber, enabling manipulation and monitoring of transfection over several days.

### 3.1. Mask and wafer fabrication

- All mask and wafer fabrication steps are performed in a class 100 clean room. For researchers without access to a clean room or PDMS fabrication facilities, many commercial mold and device fabrication services are available. The design files of the transfection device are available for download from zenodo.org (https://zenodo.org/record/820777#.WVSrbhPfrOQ).

#### 3.1.1. Chip design

1. The microfluidic device is designed using CAD software. The chip measures 1.6 x 5.8 cm and contains 280 culturing and transfection chambers (Fig. 2a). Two molds should be designed: one for the control layer (30 μm height), which contains the valves, and another for the flow layer (30 μm height), which contains the channels and chambers necessary for reagent introduction and cell culturing (Fig. 2b,c, Note 3).
2. The flow layer mold contains two patterns (SU-8 and AZ9260) on the same wafer, so alignment marks are added to the design to enable alignment of the layers during the exposure step.
3. The control layer is scaled by 101.5% to account for PDMS shrinkage during curing.

**Figure 2.**
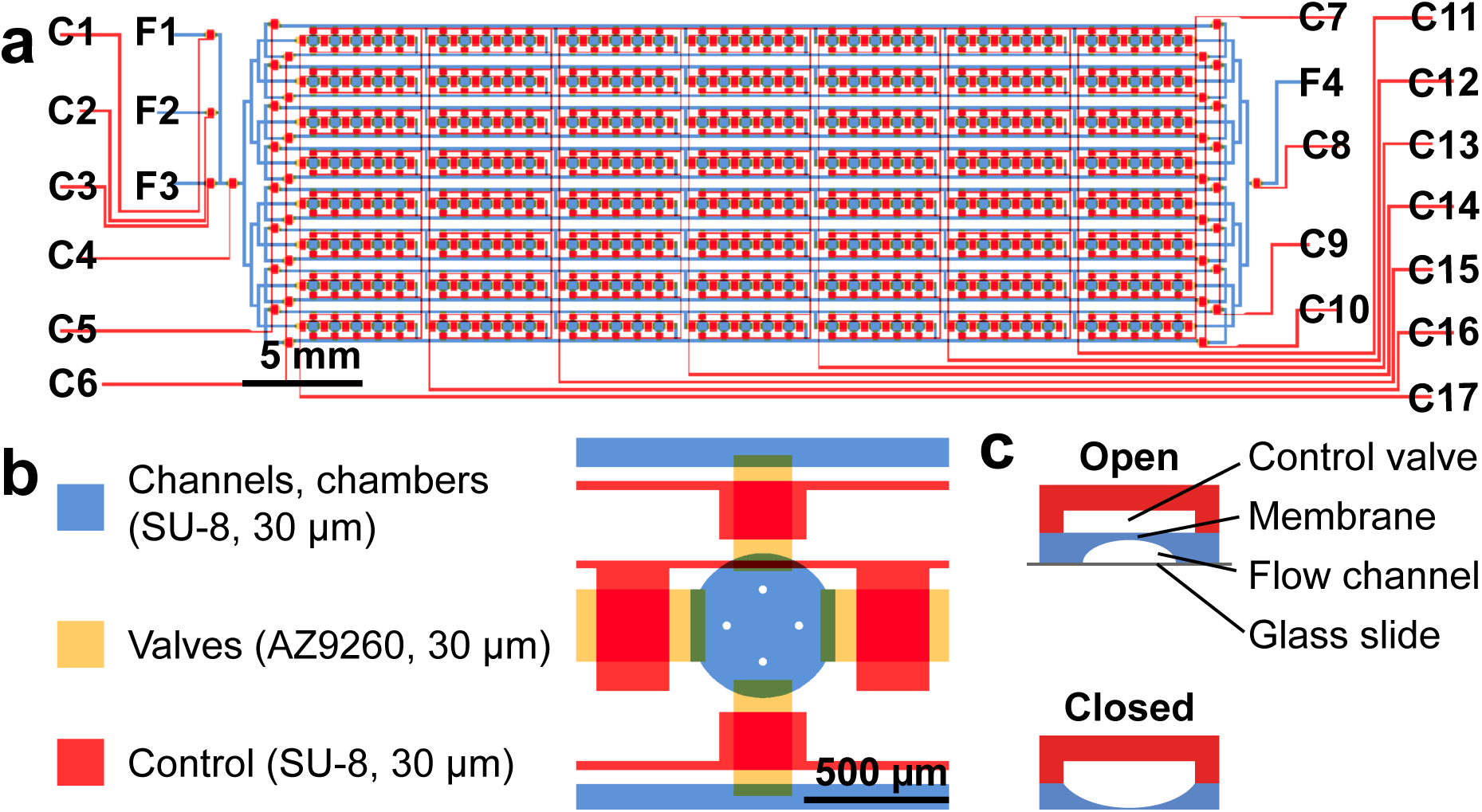
Design of the high-throughput transfection chip. Design of the 280-chamber transfection and cell culturing chip. Red, control lines for valve actuation, blue/yellow: flow channels and cell culturing chambers. F1-F3 are flow inputs, F4 is the flow outlet. C1-C3 control inputs F1-F3. C4 and C8 control access to the body of the chip. C7 controls access of cell chambers to medium channels. C5 and C9 control flow in the upper medium channel. C6 and C10 control flow in the lower medium channel. C11-C17 segregate the chambers in individual columns. Close-up of one chamber. The features are created by patterning different layers of photoresist. (c) Schematic of valve opening and closing. When the control line is actuated, the resulting pressure depresses the PDMS membrane separating the control valve and the flow channel, closing the flow channel.

#### 3.1.2. Mask fabrication

1. A Heidelberg VPG200 laser lithography system with a 20 mm writing head and a UV-light source (i-line λ = 355 nm) is used to write each layer of photoresist on a separate mask (2 masks for the flow layer, 1 for the control layer).
2. Masks are developed using the DV10 instrument. The development dispenser is purged before the mask is placed inside the chamber and developer (a 1:5 MP 351:DI water mixture) is applied twice (15 s application followed by 45 s agitation), followed by rinsing and drying.
3. The chrome layer of the mask is etched in a perchloric acid solution for 120 s, then washed with water (quick dump rinse followed by an ultra pure water bath) and dried. The remaining photoresist is removed using TechniStrip 1316 (manual application to mask followed by complete immersion in bath for 10 min). The mask is washed with water (quick dump rinse followed by an ultra pure water bath) and air dried.

#### 3.1.3. Control layer wafer fabrication

1. Wafers are cleaned with oxygen plasma for 10 min at 500 W power, 2.45 GHz plasma frequency, Gas: O_2_ and CF_4_, max. flow 500 ml/min.
2. Immediately after, the Sawatec LSM250 is used to spin coat a 30 μm thick layer of SU-8 (GM1070): 5 s ramp up, 5 s at 500 rpm, 20.3 s ramp up, 40 s at 2527 rpm, 1 s at 3527 rpm, 1 s ramp down to 2527 rpm, 5 s at 2527 rpm, and 25.3 s ramp down to 0 rpm.
3. The wafers are relaxed for 20-45 min at room temperature.
4. The wafers are baked using the hot plate of the Sawatec LSM250 for 5 min at 130°C, with ramp up and ramp down times of 50 min each.
5. The MA6 mask aligner is used to expose the baked wafers to light for 8 s in soft contact mode with an alignment gap of 30 μm, using a lamp intensity of 20 mW/cm^2^ (WEC type: cont, N2 purge: no, WEC-offset: off).
6. The wafers are baked on the Sawatec LSM250 hot plate as follows: 25 min ramp up, 15 min at 70°C, 15 min ramp up, 40 min at 100°C, 60 min ramp down to room temperature.
7. The wafers placed in a storage box overnight to encourage rehydration under ambient conditions.
8. The wafers are developed in 2 consecutive 5 min PGMEA baths, followed by 1 min in isopropanol and air drying. If white residues appear during the isopropanol bath, the wafer is returned to the PGMEA bath for further development.
9. The wafers are inspected by microscopy, and if cracks are visible in the features, a hard bake (120 min at 135°C, with ramp up and down times of 40 min each) is performed.
10. The features are measured using a Dektak XT surface profilometer.

#### 3.1.4 Flow layer wafer fabrication

1. The flow layer is patterned with SU-8 (for the channel features) as described above. For the valve features, AZ9260 is next patterned on the same wafers as described below.
2. An automated wafer coating, baking, and development machine (EVG150) is used. The wafers are dehydrated for 4 min 30 s at 160°C (see Note 4). AZ9260 is then spin coated as follows: dispensing of resist at 100 rpm, increase to 500 rpm, 3 s ramp up, 45 s at 1700 rpm, 60 s ramp down to 0 rpm. Edge bead removal (15 s) and backside removal (20 s) are performed at 1000 rpm. The coated wafers are baked for 1 min 35 s at 110°C. AZ9260 resist is deposited a second time (using the same recipe) to yield a final height of 30 μm. The wafers are baked for 3 min 15 s at 110°C.
3. The coated wafers are rehydrated overnight.
4. The MA6 mask aligner is used to expose the wafers to light for 2 sessions of 28 s, separated by a 10 s wait. The mask used with the exposure tool is aligned to the existing SU-8 pattern on the wafer. The exposure tool is operated in hard contact mode with an alignment gap of 30 μm and a lamp intensity of 20 mW/cm^2^ (WEC type: cont, N2 purge: no, WEC-offset: off).
5. The EVG150 is used to dispense developer AZ 400K onto the wafers (9 min at 250 rpm), rinse them with DI water (30 s at 250 rpm), perform backside removal (10 s at 250 rpm), and drying (30 s at 3000 rpm).
6. The wafers are washed with water (quick dump rinse followed by an ultra pure water bath) to remove any remaining developer residues.
7. The features are annealed (to create a rounded profile) by placing the wafers on a Sawatec LSM250 hot plate for 300-600 s at 140°C.
8. The features are measured with a Dektak XT surface profilometer. If the profile of the valve regions is not completely rounded, the annealing bake should be repeated for a longer amount of time.

### 3.2. PDMS chip fabrication

1. Flow and control wafers are incubated for 10 min in a sealable wafer holder containing a plastic cap filled with 1 ml TMCS. This process is repeated prior to each chip fabrication to prevent PDMS from lifting photoresist features off the wafer.
2. Both wafers are placed into glass petri dishes lined with aluminum foil (see Note 5).
3. For the control layer, 60 g of a 5:1 PDMS mixture (50 g Part A: 10 g Part B) is placed in a disposable PP cup, mixed for 1 min at 2,000 rpm (∼400 x g), and degassed for 2 min at 2,200 rpm (∼440 x g) in a centrifugal mixer.
4. The control wafer is cleaned with a nitrogen air gun to remove any debris or small dust particles (see Note 6). The PDMS mixture is immediately poured onto the control layer wafer and the assembly is degassed in a vacuum chamber for 20 min. Upon removal of the PDMS-coated wafer from the chamber, any remaining bubbles are pushed to the edges of the petri dish using a pipet tip.
5. For the flow layer, 20 g of a 20:1 PDMS mixture (20 g Part A: 1 g Part B) is placed in a disposable PP cup and mixed and degassed as described above.
6. The flow wafer is placed inside the spin coater and cleaned with a nitrogen air gun, then coated with PDMS using a ramp of 15 s and a spin of 35 s at 650 rpm.
7. Both the control and flow molds are baked in an oven for 30 min at 80°C (see Note 7).
8. The molds are removed from the oven and the PDMS of the control layer is cut with a scalpel to remove each chip design from the wafer. To enable attachment of tubing to the control channels, holes are punched at the control line inlets. The holes are punched with the patterned side of the chip facing up, and displaced PDMS stubs are removed with tweezers.
9. Scotch tape is placed in contact with the edges and patterned side of the chip and subsequently removed to clean debris from the surface.
10. The control layer chips are aligned to the PDMS-coated flow wafer using a stereomicroscope. Slight pressure is applied throughout the chip to ensure that all regions of the control layer are in contact with the flow layer.
11. The assembly is baked for 90 min at 80°C.
12. A scalpel is traced around the outline of each device to cut the PDMS layer coating the flow wafer (see Note 8).
13. The bonded control/flow devices are removed from the wafer, and holes for the flow inlets and outlets are punched with the patterned side of the chip facing up.
14. The edges and patterned side of the chip are cleaned with Scotch tape before bonding to the transfection arrays.
15. For the flow wafer, the remaining thin PDMS layer is removed by pouring a 10:1 PDMS mixture (mixed and degassed) onto the wafer and baking for 30 min at 80°C. Once cured, the thick PDMS layer can be easily removed from the entire wafer.
16. For the control wafer, excess PDMS is cut away from the body of the wafer and a frame is left near the edges to reduce the quantity of PDMS required for subsequent batches of chips.

### 3.3. PLL microarraying

1. Glass slides are arranged in a rack and submerged in a covered dish containing a 57% Ethanol, 10% w/v NaOH solution. The slides are incubated for 2 h with gentle shaking, then rinsed 5x with Milli-Q water and air dried.
2. The arraying pin is placed in a 15-ml conical tube containing a soap-water mixture (∼0.5 ml dish soap diluted in 15 ml water) and placed in a sonication bath for 10 min (see Note 9). The soap-water mixture is then replaced with 70% ethanol and the pin is sonicated for another 10 min. The pin is dried with a nitrogen air gun and placed in the microarrayer head.
3. The PLL sample is prepared by adding 25 μl of a 0.1% PLL solution to 50 μl of a 0.225 M, pH 8.4 boric acid solution. The sample is loaded into a well on a 384-well plate. One sample can be used to prepare up to 2 glass slides. To simultaneously prepare more glass slides, multiple PLL samples should be used.
4. The glass slides and the well plate are loaded into the microarrayer (see Note 10). PLL is stamped 4 times per spot in a cyclic fashion, with approximately 8 min between cycles (modifying the wait times to 6-14 min has no detrimental effect on transfection efficiency). The pitch of the spots matches that of the chambers on the microfluidic chip: 8 rows, 41 columns (7 sets of 5, with one blank position between each set), x-pitch 900 μm, y-pitch 1700 μm. Spotting parameters are 1 s inking time, 500 ms printing time, 4 stamps/ink. Every 40 inks, the pin is washed with water for 500 ms and dried for 500 ms. Humidity in the spotter is set to 50%.
5. Orientation marks are placed on the slides (see Note 11) and 2 h after arraying is complete, PLL arrays are washed 5x 30 s with Milli-Q water, dried using a nitrogen air gun, and stored in a desiccator. PLL slides should be used between 2 and 8 weeks post-coating, since the quality of the PLL has been shown to deteriorate beyond this time period [20].

### 3.4. Lipid-DNA microarraying

For a modification of this method (gelatin-DNA method), see Note 12.

1. A 0.5% gelatin solution is prepared as follows: 0.15 g of gelatin is added to 30 ml of Milli-Q water that is already at 60°C. The mixture is incubated in a 60°C water bath and swirled gently 8x during a 15 min period. The solution is removed from the bath and left at room temperature until it has cooled to 37-40°C. The mixture is then loaded into a syringe, filtered through a 0.45 μm membrane, and stored in 700 μl aliquots at 4°C (see Note 13).
2. Sucrose is dissolved in EC buffer (Effectene transfection kit) to a final concentration of 0.2 M, and the solution is filtered.
3. 1.5 μg of supercoiled plasmid DNA is diluted in 15 μl of the 0.2 M sucrose-EC buffer (see Note 14). In the case of co-transfection the total DNA of all plasmids should add up to 1.5 μg and the mixture should be vortexed for 10 s, then incubated for 15 min. The tubes are quickly spun down after each vortexing step.
4. 1.5 μl Enhancer (Effectene transfection kit) is added. The mixture is vortexed for 1 s and incubated for 5 min.
5. 5 μl Effectene is added. The mixture is gently vortexed for 10 s and incubated for 10 min.
6. 12.7 μl of the 0.5% gelatin solution and 12.7 μl of a 0.1% fibronectin solution [12] are added and mixed by pipetting up and down.
7. Samples are transferred to a 384-well plate for microarraying. Each sample can be used to array up to 140 spots.
8. The microarray pin is cleaned as previously described and placed in the microarray head. The PLL-patterned slides are oriented in the holders and the spotter is programmed to spot in the same positions as previously used for PLL.
9. The lipid-DNA is stamped once per spot (see Note 15). Spotting parameters are 1 s inking time, 500 ms printing time, 4 stamps/ink. Every 40 inks, the pin is washed with water for 500 ms and dried for 500 ms. Humidity in the spotter is set to 65%.
10. Printed slides are stored in a desiccator and used within a few weeks.

### 3.5. Device assembly

1. The spotted glass slides are cleaned with a nitrogen air gun and placed into a plasma chamber with the spots facing up.
2. The PDMS chips are cleaned with Scotch tape and placed into the plasma chamber with the features facing up.
3. The slides and chips are simultaneously plasma treated (settings: 23 ml oxygen, power setting of 5, 1 min vacuum pump, 1 min gas, 7 s plasma). Immediately afterwards, the functionalized surfaces are aligned (one-shot) using a stereomicroscope (see Note 16). Slight pressure can be applied to ensure that all parts of the chip are in contact with the glass slide (see Note 17). The assembly is baked for 1 h at 80°C.
4. The transfection devices are stored in a desiccator.

### 3.6. Microfluidic setup and operation

The microfluidic setup comprises pressurized manifolds for flow and control layer manipulation, a fluorescence microscope, an incubation chamber, and a camera (Fig. 3).

**Figure 3.**
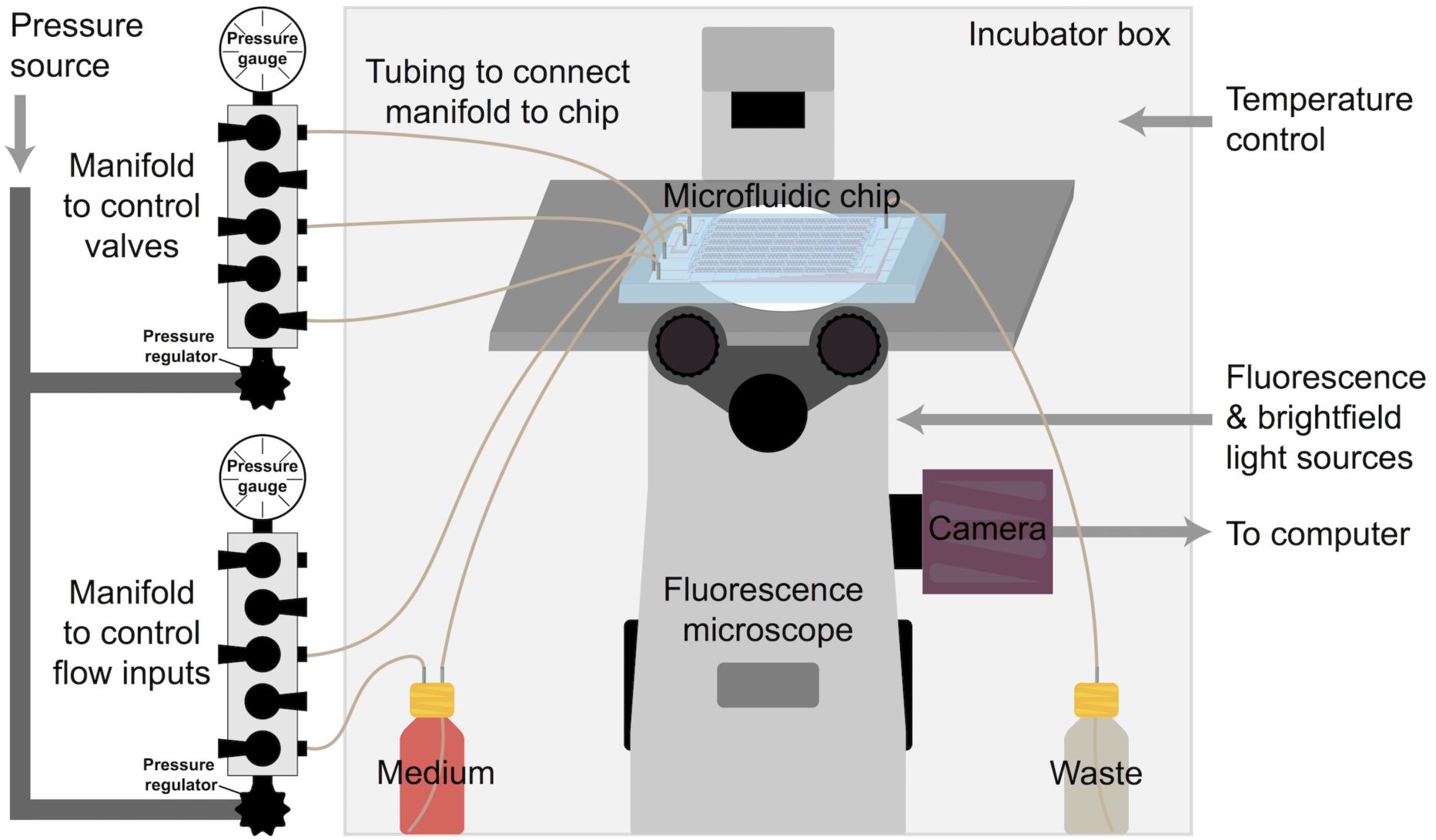
Schematic of the microfluidic setup. A pressure source is connected to manifolds that enable manipulation of specific control and flow lines on chip. The ports of the manifold are individually controllable and attached to the chip with flexible tubing. The chip is cultured on the stage of a fluorescence microscope contained inside an incubation chamber.

#### 3.6.1. Microfluidic setup: control layer

1. A pressure source is attached to manifolds controlled by pressure regulators, with gauges connected to the ends of the manifolds. The toggle switches on the manifolds control the on/off state of individual ports. Each port is connected to a flexible Tygon tube that is fitted with a steel pin at the other end, enabling connection to the holes punched on the chip. The pneumatic setup can be custom built to allow for different pressure ranges and to accommodate larger numbers of control and flow lines.
2. A syringe is used to aspirate and transfer filtered water to a control line tube (a short piece of Tygon tubing serves as an adaptor between the syringe needle and the pin). The control line is placed in the corresponding hole of the chip and pressurized at 5 psi (see Note 18). This process is repeated for each control line (Fig. 1a, C1-C17). Once the air in the chip’s control lines has been replaced by water, the pressure is increased to 22 psi.

#### 3.6.2. Microfluidic setup: flow layer

1. For flow connections, small samples (such as PBS for washing or cells for loading) are contained in Tygon tubing attached to a manifold at one end and the chip at the other end. The Tygon tubes are loaded and connected as described above for the control lines.
2. For large volume or long-term flow connections, PTFE tubing is used (see Note 19). A bottle containing filtered CO_2_-independent medium supplemented with 10% FBS, antibiotic-antimycotic, and glutamax is attached to the microfluidic setup (see Note 20). A piece of tubing is placed inside the bottle, running from top to bottom. The tube is connected to a pin that punctures the septa of the cap (this entire assembly is autoclaved prior to use). On the other side of the septa, the pin is connected to a longer piece of tubing that connects to the chip. Finally, a second piece of tubing is attached to a manifold at one end and punctures the septa on the other end. This setup pressurizes the bottle to drive flow into the chip (Fig. 3).

#### 3.6.3. Microfluidic operation: flow inputs

1. To flow samples on chip, all valves are initially closed (control lines C1-C17 pressurized).
2. The tube containing the sample is connected to the chip (for example to position F1) and pressurized at ∼2-4 psi.
3. The corresponding valve (C1) controlling the flow of F1 is opened.
4. A second valve (such as C3) controlling an additional flow input (such as F3) is opened, serving as a purge to remove any air introduced at the end of the sample.
5. To terminate the purge and begin flowing on chip, C3 is activated and the master valves (C4, C8) controlling access to the body of the chip are opened. The outlet, F4, is connected to a waste bottle with Tygon tubing.
6. All valves controlling channel and chamber features (C5-C7, C9-C17) are opened to facilitate filling of the entire chip.
7. To disperse any air bubbles trapped in the chip, the outlet valve (C8) is activated while the sample is flowed into the chip. The pressure can be increased to accelerate this step. Once all air has been removed, C8 is opened to permit continuous flow.

#### 3.6.4. Microfluidic operation: cell loading

1. A T-75 flask of HEK 293-T cells at 70-80% confluence is harvested. 800,000 cells are centrifuged and resuspended in 20 μl PBS (see Note 21).
2. As quickly as possible, the sample is loaded into a Tygon tube, attached to the manifold and chip, and purged as described above.
3. The cells are loaded into the chip at a rate of 27 μl/min (see Note 22), with columns loaded in sets sequentially (one column of 5-chamber segments at a time, Fig. 4a). When a sufficient number of cells (∼70% confluent) is obtained in the chamber segment (see Note 23), chamber-separating valves (C11-C17) are immediately actuated to capture cells in the chambers (see Note 24).

**Figure 4.**
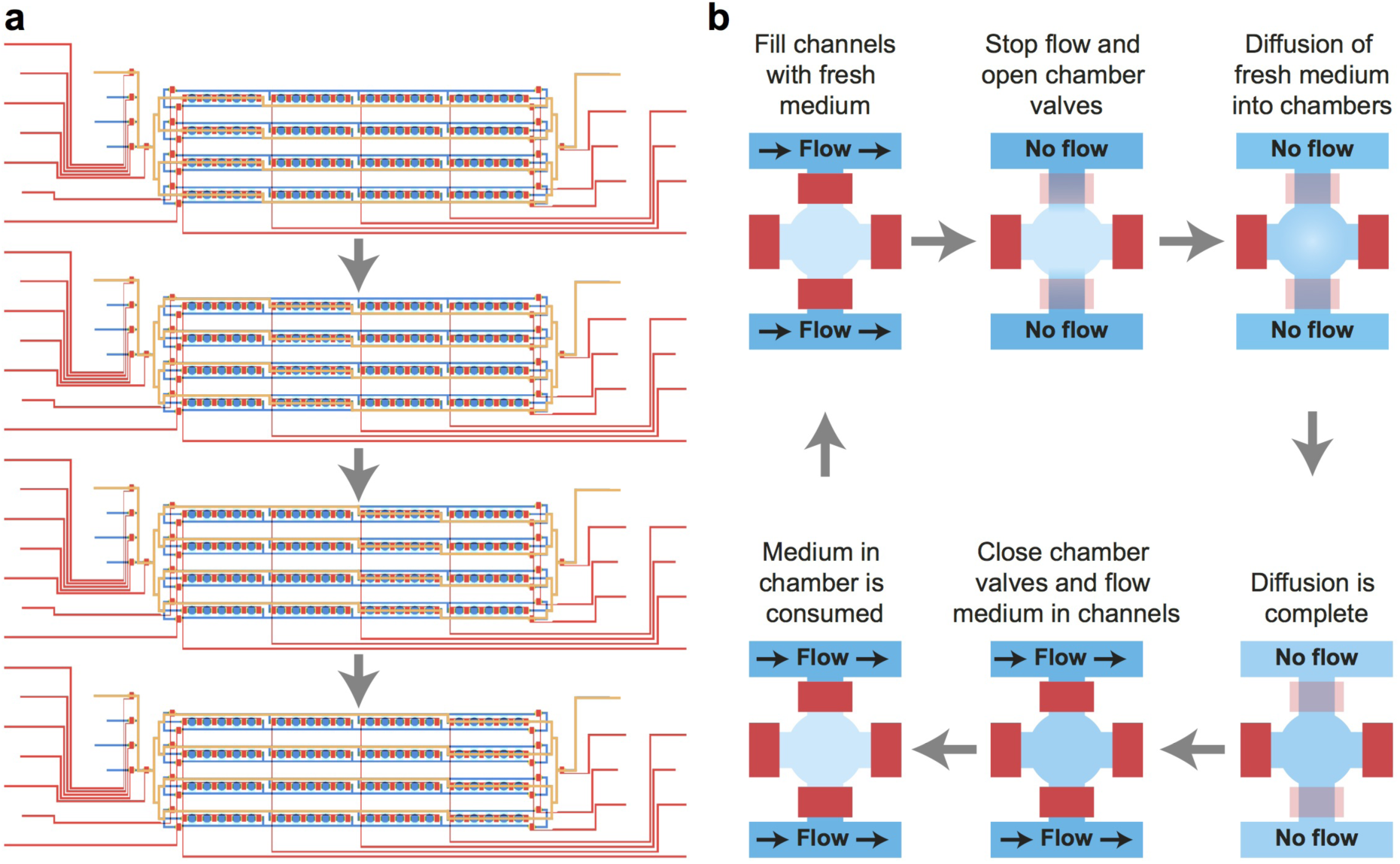
Schematic of cell loading and medium perfusion. (a) Cell loading is performed by sequentially loading each segment of 5 chambers. The yellow lines designate the flow trajectory through the chip. A low-throughput chip (4 columns) is shown for clarity, but the procedure is the same for the 7-segment high-throughput chip. (b) Medium is pulse perfused for the first hour after cell loading to promote attachment to the surface. The device is switched between 2 states: (1) flow of medium through the channels with access to the cell chambers closed, and (2) no flow, with exposure of chambers to the channels for medium diffusion.

#### 3.6.5. Microfluidic operation: cell culturing

1. The chip is cultured on the stage of an automated microscope contained in an incubation chamber maintained at 37°C.
2. For the first hour after loading, medium is pulse perfused [21] to promote attachment and prevent cells from being washed out of the chambers (Fig. 4b, Note 25). Medium is flowed for 5 min through the channels with access to the chambers closed. For the next 5 min, the flow of medium is stopped and the valve joining the chambers and medium channels are opened.
3. After 1 h of pulse perfusion, the top/bottom chamber valves are opened and medium is continuously flowed at a rate of ∼1 ml/h (see Note 26).

### 3.7. Imaging

1. To measure transfection efficiency, images are acquired 48 h (see Note 27) after introducing cells onto the chip, using 20x magnification to capture an entire chamber in the field of view (Fig. 5a). Images can also be acquired continuously after cell loading to monitor the progression of protein expression. Chambers are imaged in fluorescence and brightfield modes to count both the number of fluorescent cells and the number of total cells.
2. Images can be stitched together (for example using the Grid/Collection Stitching plugin in Fiji [22]) to reconstruct the entire array (Fig. 5b). Cells can be counted using programs such as CellProfiler or Matlab.
3. Transfection efficiency is calculated as the area occupied by fluorescent cells in the entire chamber (500 μm diameter) divided by the area occupied by cells within the 300 μm diameter spot where the lipid-DNA was deposited (see Note 28).

**Figure 5.**
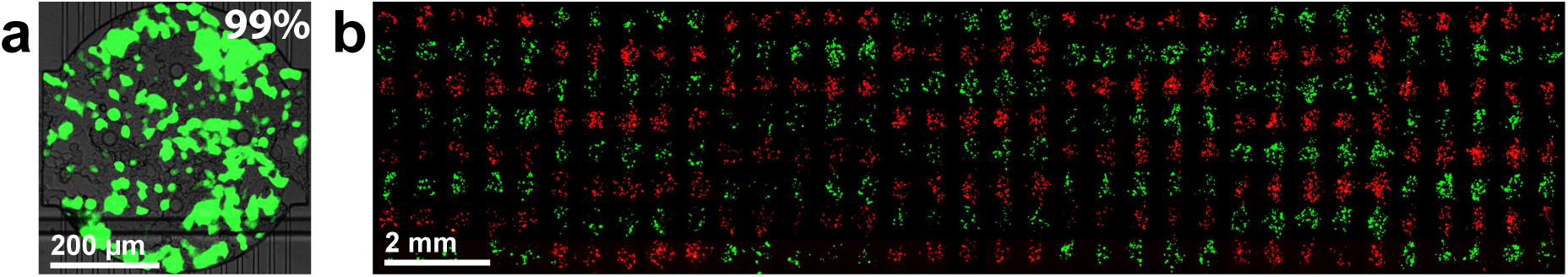
Transfection on the high-throughput chip. (a) Composite fluorescence image showing high efficiency eGFP transfection on the device. (b) Fluorescence micrograph of an entire 280-chamber chip, with each 5-chamber segment transfected with a plasmid for expression of eGFP (green) or tdTomato (red).

## NOTES

1. DNA samples should be supercoiled plasmids obtained using standard plasmid preparation kits, with a concentration of 200-500 ng/μl to avoid significant dilution of the transfection mixture.
2. The FBS should be of high quality and subjected to limited freeze-thaw cycles to ensure high transfection efficiency.
3. Designs intended for use with positive photoresist should be inverted before mask writing to ensure proper polarity, and a frame should be added around the features to provide sufficient empty space around the design.
4. If problems with AZ9260 adhesion to the wafer are experienced, an HMDS bake may be performed prior to dehydration and coating.
5. Multiple layers of foil may be necessary to prevent holes. The foil prevents direct contact and bonding between the PDMS poured on the wafer and the glass dish.
6. To fabricate functional microfluidic chips, it is essential to work quickly, protect the exposed flow layer, and keep wafers and chips free of dust. Chip fabrication can alternatively be performed in a clean room environment. Wafers should never be cleaned with water or organic solvents.
7. Wafer-containing glass petri dishes should not be stacked in the oven, since this results in sub-optimal heat distribution.
8. Space allowing a strip of PDMS can be removed along each of the 4 edges of the chip to ensure a clean cut of both the flow and control layers. Small diagonal cuts made at each corner will also discourage delamination during chip operation.
9. Care should be exercised when handling microarraying pins. The tips of the pins are easily damaged, which results in printing or alignment problems.
10. Glass slides should be carefully aligned to the bottom right corner of the microarray holder positions, and vacuum should be applied to prevent the slides from moving. Proper alignment is critical since a second round of spotting (lipid-DNA) will be performed on the same slides and the lipid-DNA must be deposited precisely on top of the PLL spots.
11. PLL spots are no longer visible after the slides have been washed to remove excess PLL. To enable proper orientation of the slide in the holder for subsequent lipid-DNA spotting, the bottom right corner of the PLL slide is marked with a permanent marker prior to washing.
12. For the gelatin-DNA technique, DNA mixtures lacking transfection reagent are spotted and the Effectene is flowed on chip prior to cell loading. Samples are prepared similarly to the lipid-DNA mixtures and contain 1.5 μg DNA and buffer EC (total of 25.4 μl), 12.7 μl gelatin, and 12.7 μl fibronectin. A mixture of 16 μl Enhancer, 150 μl EC buffer, and 25 μl Effectene is flowed on chip for 30 min using the same conditions used for medium perfusion. Medium is flowed on chip for 20 min prior to cell loading.
13. Proper gelatin preparation is important for transfection efficiency, and aliquots should not be stored longer than 1 month.
14. In the case of limited DNA samples, the lipid-DNA recipe can also be halved.
15. Depositing lipid-DNA multiple times per spot does not increase transfection efficiency, and as has been previously reported [17] results in imprecisely localized spots due to spreading of excess DNA.
16. A maximum of 2 chips should be plasma treated at a time since alignment is time sensitive and must be performed immediately after plasma treatment.
17. Application of high pressure to the chip immediately after plasma treatment should be avoided since valves or chambers may depress, resulting in permanently closed valves or collapsed chambers.
18. When filling control lines and flow samples, the end of the tube fitted with the pin should be positioned downward and the free end (for connection to the manifold) should be elevated. In addition to careful aspiration with the syringe, this configuration discourages the sample from sliding further back into the tube, minimizing bubbles introduced on chip.
19. PTFE tubing is used to supply long-term culturing medium because cell toxicity may be observed when using Tygon tubing, as has been previously reported [23].
20. The medium bottle should be carefully handled since any frothing may introduce bubbles onto the chip.
21. As with traditional transfection, efficiency is increased when using low passage number cells. Transfection of different cell types (e.g. CHO) is possible, but the composition of the lipid-DNA mixture may need to be optimized accordingly.
22. Flow rate is determined by measuring the volume of liquid exiting the chip over a period of time.
23. In the case of clogging during cell loading, additional pressure may be applied to disperse the clog. To ensure homogeneity, it is recommended to purge a large amount of the sample (discard 10-50%) before running the cells through the chip for loading.
24. Example of chip loading (left to right) for a cell sample connected to F1. Cells will enter the first column through the middle channel and exit through the bottom right (all valves closed except C1, C4, C8, C10 and C17). For columns 2-6, cells enter through the top left and exit through the bottom right (all valves closed except C1, C4, C5, C8, C10, and C12-C16, depending on the column being loaded). For the last column, cells enter through the top left and exit through the middle channel (all valves closed except C1, C4, C5, C8, and C11).
25. Example of pulse perfusion from F2, medium flow stage: all valves closed except C2, C4-C6, C8-C10. Medium diffusion stage: all valves closed except C2, C5, C6, C7, C9, C10. To automate pulse perfusion, solenoid pneumatic valves arranged on a manifold (Pneumadyne) and controlled by a custom written LabVIEW VI program can be used.
26. Example of continuous medium flow from F2: all valves closed except C2, C4-C10). Long-term (longer than 48 h) culturing on the chip may result in cell outgrowth from the chamber and migration into the medium perfusion channels, causing clogging and preventing proper medium flow. To discourage this, the C7 valve can be periodically opened and closed to crush and disperse outgrown cells. Another concern is the presence of bubbles in the chip during culturing, which may be mitigated by closing the outlet (C8) valve while the chip is pressurized with medium. In the case of frequent problems with bubbles, an automated lab program such as LabVIEW may be used to periodically close C8.
27. For co-transfection experiments, imaging at time points beyond 48 h may be necessary (for example, for long maturation times, or for co-transfections in which one protein must first be expressed before activating the expression of a second protein).
28. Normalization when calculating transfection efficiency is necessary because only some of the cells loaded into the chamber have access to the transfection mixture (a 300 μm spot within a 500 μm chamber). Some cells that initially settle in the lipid-DNA area migrate to different parts of the chamber after 48 h (when images are captured), resulting in the dispersed pattern visible in the images.

